# Rapid selection response to ethanol in *S. eubayanus* emulates the domestication process under brewing conditions

**DOI:** 10.1101/2020.12.07.415240

**Authors:** Wladimir Mardones, Carlos A. Villarroel, Valentina Abarca, Kamila Urbina, Tomás A. Peña, Jennifer Molinet, Roberto F. Nespolo, Francisco A. Cubillos

## Abstract

Although the typical genomic and phenotypic changes that characterize the evolution of organisms under the human domestication syndrome represent textbook examples of rapid evolution, the molecular processes that underpin such changes are still poorly understood. Domesticated yeasts for brewing, where short generation times and large phenotypic and genomic plasticity were attained in a few generations under selection, are prime examples. To experimentally emulate the lager yeast domestication process, we created a genetically complex (panmictic) artificial population of multiple *Saccharomyces eubayanus* genotypes, one of the parents of lager yeast. Then we imposed a constant selection regime under a high ethanol concentration in 10 replicated populations during 260 generations (six months) and compared them with evolved controls exposed solely to glucose. Evolved populations exhibited a selection differential of 60% in growth rate in ethanol, mostly explained by the proliferation of a single lineage (CL248.1) that competitively displaced all other clones. Interestingly, the outcome does not require the entire time course of adaptation, as four lineages monopolized the culture at generation 120. Sequencing demonstrated that *de novo* genetic variants were produced in all evolved lines, including SNPs, aneuploidies, INDELs, and translocations. In addition, the evolved populations showed correlated responses resembling the domestication syndrome: genomic rearrangements, faster fermentation rates, lower production of phenolic-off flavors and lower volatile compound complexity. Expression profiling in beer wort revealed altered expression levels of genes related to methionine metabolism, flocculation, stress tolerance and diauxic shift, likely contributing to higher ethanol and fermentation stress tolerance in the evolved populations. Our study shows that experimental evolution can rebuild the brewing domestication process in “fast motion” in wild yeast, and also provides a powerful tool for studying the genetics of the adaptation process in complex populations.

## INTRODUCTION

Living organisms are continually adapting to changing environments by natural selection, latently harboring the raw genetic variation required for such responses. When new conditions arise, adaptation to almost every environmental scenario is possible (e.g., temperature, oxygen and nutrients) [1, 2]. In this context, the genomic analysis of human-made populations (i.e., population genomics of domesticated species) is a relatively new matter, and constitutes a promising research approach for the experimental study of evolutionary processes [3]. Nevertheless, studies that search for the causal factors shaping the genetic structure of yeast and fungal populations, such as small nucleotide polymorphisms (SNP), insertions or deletions (INDELS), copy number variation (CNV) and structural variants (SV), are still insufficient to fully characterize the integrated adaptation process to new environments [4].

Adaptive evolution in microorganisms is a process that occurs ubiquitously, including in artificial settings where micro-environments are created, and allows the adaptation of populations to defined conditions, driving the evolution process (domestication) [5]. Domestication is a stereotyped adaptive process (a “domestication syndrome”, see [6, 7]) within a human-created environment, where several characteristics can be tracked and defined as ‘domestication signatures’. These signatures are present in different fungal species, including *Aspergillus oryzae* in soy sauce [8], *Penicillium* molds associated with cheese [9] and *S. cerevisiae* [10, 11] together with *S. pastorianus* [12], responsible for beer fermentation. In this context, spore production and viability, metabolic remodeling, changes in volatile compound production, transcriptional re-wiring and faster growth rates are considered key traits and goals of microbe domestication. In the case of brewing, the yeast re-utilization process led to new spontaneous mutations generated during cell division, which, together with selective environmental pressures, such as high ethanol concentrations, selected fitter individuals [13]. Genomic analysis in beer yeast domesticated strains demonstrated the presence of common genetic patterns, such as large genomic rearrangements, aneuploidies, high heterozygosity levels and infertility, all of which are hallmarks of the adaptation process [11, 14-16].

Two main types of yeasts suffered domestication under different brewing settings; *S. cerevisiae* that ferments ale beers at temperatures near 20 °C, and *S. pastorianus* that produce lager beers fermented at lower temperatures (8-15 °C) [17]. *S. pastorianus* is an interspecific hybrid from the cross between *S. cerevisiae* and the cryotolerant wild yeast *S. eubayanus* [18]. The hybrid nature of *S. pastorianus* confers a series of competitive advantages in the fermentation environment, likely due to the combination of performance at relatively cold temperatures, efficient sugar uptake and metabolic switching between sugar sources [19]. During an intense domestication process over approximately 500 years, lager beers have evolved reduced organoleptic complexity, mainly characterized by the presence of ester compounds and the absence of phenolic off-flavors [20]. This is reflected in the absence of *PAD1* and *FDC1* in *S. pastorianus,* genes which are responsible for the synthesis of such off-flavors [21, 22] and present in *S. eubayanus*. Lager yeast domestication is characterized by a reduced lag phase in the switch from glucose to maltose, and regulatory cross-talk between *S. cerevisiae* and *S. eubayanus* sub-genomes, which complement each other in terms of the genes required for maltose/maltotriose metabolism [23, 24].

Given the recent discovery of *S. eubayanus,* its puzzling origin and apparently co-evolutionary association with *Nothofagus* trees, several authors have analyzed the worldwide distribution of *S. eubayanus*, together with its genetic, phenotypic, and fermentative diversity [24–27]. Patagonian isolates of *S. eubayanus* exhibit the most extensive genetic diversity, and the presence of the most significant number of lineages compared to Northern hemisphere populations, including five different lineages and a large group of admixed isolates [26, 27]. To date, there is no evidence of *S. eubayanus* isolates in Europe, where the original *S. pastorianus* hybrid likely originated. Interestingly, fermentation capacity varies significantly between *S. eubayanus* isolates, possibly due to differences in maltose consumption and diauxic shift capacity, resulting in two opposite outcomes: successful or stuck fermentations [27]. These isolates produce fruit and floral flavors in beer [28], but high levels of 4-vinyl guaiacol, considered a phenolic off-flavor that provides a clove-like aroma, which is not preferred among consumers [21, 28, 29] [30].

Although different reports have provided insights into the genomic and phenotypic changes responsible for the brewing capacity of *S. pastorianus,* particularly the *S. cerevisiae* genome portion, we know little about the process of *S. eubayanus* domestication before or after hybridization. Thus, further evidence is needed to understand the molecular mechanisms underpinning the *S. eubayanus* fermentative phenotype, which in turn will provide important insights into the inherent evolutionary process represented by directional selection for domestication, and correlated responses. In this study, a genetically complex artificial mixture of 30 different genotypes of *S. eubayanus* was continually exposed to high ethanol levels, mimicking the domestication process in breweries. We measured their correlated responses including their genomic, transcriptomic and phenotypic changes, and identified candidate genes that confer ethanol tolerance. Our results demonstrate that a single genetic background consistently overcomes the remaining strains, showing greater fermentation performance, but also significantly higher fitness in oxidative and osmotic stress environments. To an extent, we thus recreate the domestication process in the laboratory, showing how this cryotolerant yeast adapted to the competitive beer environment of a human industry and proved that experimental evolution can rebuild the brewing domestication process in *S. eubayanus* in “fast motion”. This provides a powerful tool for disentangling the molecular, physiological and biochemical processes that underlie the domestication of domesticated microorganisms.

## MATERIALS AND METHODS

### Microorganisms and culture media

Thirty *S. eubayanus* strains isolated from bark samples obtained from *Nothofagus pumilio* trees in south Chile were utilized for the experimental evolution assay, as listed in **Table S1**. These strains were previously reported and belong to the Patagonia B cluster [27]. *S. cerevisiae* L299 [31] and MTF2444 (EC1118 *hsp12::GFP)* [32] strains were used as growth control and in the competition assays, respectively. Additionally, we used the *S. pastorianus* Saflager W-34/70 (Fermentis, France) strain as a lager fermentation control. All isolates were maintained in YPD agar media (yeast extract 1%, peptone 2%, glucose 2% and agar 2%) and stored at −80°C in 20% glycerol stocks.

### Experimental evolution

Initially, one colony from each *S. eubayanus* strain was cultured in 0.67% yeast nitrogen base (YNB) media (Difco, France) with 2% glucose at 20°C (hereinafter referred to as GLU) and 150 rpm orbital shaking. Later, each pre-inoculum was utilized to prepare a co-culture in a single 250 mL flask to obtain a final concentration of 1×10^6^ cells/mL of each strain. Ten replicates were set up (parallel populations) in 5 mL GLU and ten supplemented with 0.67% YNB media, 2% glucose and 9% ethanol (hereinafter referred to as EtOH). The inoculum was resuspended and transferred to the 20 replicates to obtain a final concentration of 1×10^6^ cells/mL (**Figure 1A**). The adaptative evolution assays were performed at 20°C at 150 rpm for 72 h. Subsequently, the cultures were used to inoculate fresh 5 mL cultures at an inoculum density of 1 × 10^6^ cell/mL, and this procedure was sequentially repeated. The number of generations was estimated using the ‘‘generations = log (final cells - log initial cells)/log2” formula, summing up the number of cells/mL doublings between every culture transfer during the adaptive evolution process.

**Figure 1.**
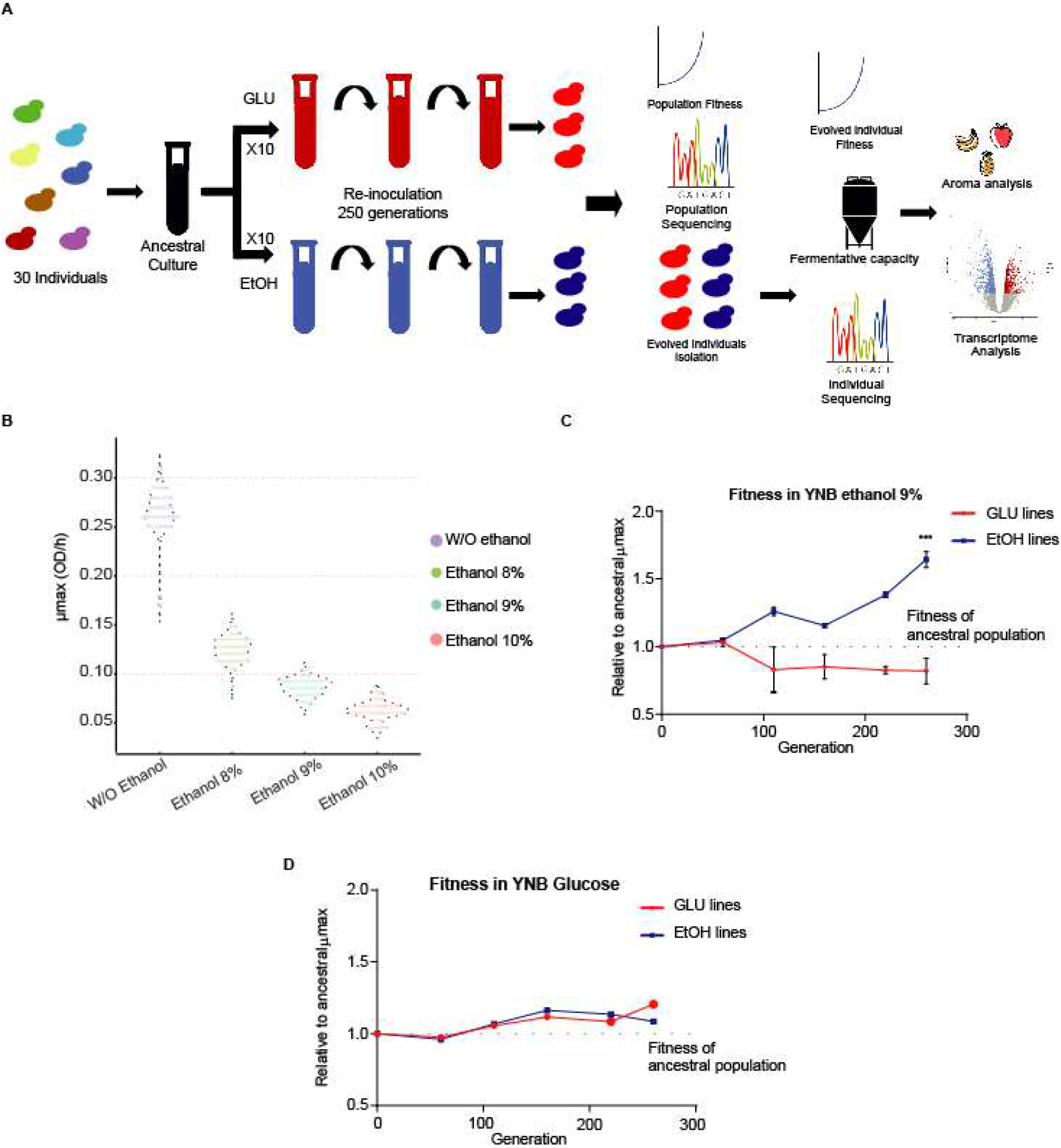
Fitness of the individual and evolved lines under ethanol. (A) Experimental evolution strategy in 10 replicated lines under YNB + glucose (GLU, red tubes) and YNB + GLU + ethanol 9% (EtOH, blue tubes). From every line, individuals were isolated and subjected to phenotyping, fermentation and sequencing analysis. (B) The growth rate (μmax) of the different parental strains used in this study was estimated under ethanol 8%, 9% and 10%. The fitness of the evolved lines under (C) ethanol and (D) glucose.

### Phenotyping assay

The phenotyping assay was performed as previously described [27]. Briefly, isolates were pre-cultivated in 200 μL 0.67% YNB medium supplemented with glucose 2% for 48 h at 25°C. Next, strains were inoculated to an optical density (OD) of 0.03-0.1 (wavelength 630 nm) in 200 μL growth media, where the following carbon sources were considered: Glucose 2%, Fructose 2%, Maltose 2%, Galactose 2%, Pilsner Beer Wort 12 °Plato (°P) and incubated without agitation at 20°C for 24 h using a Tecan Sunrise absorbance microplate reader. Additionally, several environmental stressors were assessed, including ethanol 9%, Sorbitol 20%, H2O2 3 mM, SDS 0.001% and high temperature (28 and 34°C) during 48 h. For ethanol 9%, experiments were carried out for 96 h. The OD was measured every 30 minutes using a 630 nm filter. Each experiment was performed in triplicate. Maximum growth rate, lag time and OD max parameters were obtained for each strain using the GrowthRates software with default parameters [33].

Growth curves incorporating carbon source switching from glucose to maltose and galactose were determined under micro-cultivation conditions in YP (1% yeast extract, 2% peptone) media including either 5% glucose, 5% maltose or 5% galactose at 25°C for 48 h. Precultures were grown in YP with 5% glucose medium at 25°C for 24 h. Cultures were then diluted to an initial OD600nm of 0.1 in fresh YP 5% glucose medium for an extra overnight growth. The next day, cultures were used to inoculate a 96-well plate with a final volume of 200 μL YP with the disaccharide source at an initial OD600nm of 0.1. The growth curves were monitored by measuring the OD600nm every 30 min as previously mentioned. All experiments were performed in triplicate. Lag phase and maximum specific growth rate (μmax) were estimated as previously described [34] using the R software version 3.6.3.

### Fermentations in beer wort

Fermentations were carried out as previously described [28, 29]. Briefly, fermentations were performed in at least three biological replicates, depending on the experiment, in 12 °P using a BrewFerm Pilsner commercial beer kit (Beringen, Belgium). For this, a colony was transferred to 5 mL 6 °P pilsner beer wort supplemented with 0.3 ppm ZnCl2 and incubated at 20°C with orbital shaking at 150 rpm for 24 h. Then, the complete pre-inoculum was transferred to 50 mL 12 °P pilsner beer wort and incubated in similar conditions for 24 h. Cells were utilized to inoculate 50 mL fresh 12 °P pilsner beer wort to a final concentration of 1.8 × 10^7^ cell/mL. Cultures were maintained at 12°C for 14 days without agitation and weighed every day to calculate the CO_2_ released.

Larger volume fermentations for RNA extraction and metabolite production analysis were carried out in 1.5 L 12 °P beer wort for 14 days at 12°C. At the end of the fermentation, metabolites such as glucose, fructose, maltose, maltotriose, ethanol and glycerol were estimated using HPLC [27]. Volatile compounds were detected using HS-SPME-GC-MS as previously described [28].

### Competition Assays

A total of 1 × 10^6^ cells/mL of the evolved and *S. cerevisiae* MTF2444 (EC1118 *hsp12::GFP)* strains were separately pre-incubated in 5 mL YNB media supplemented with 2% glucose for 24 h. Evolved individuals were mixed in equal proportions with the *S. cerevisiae*MTF2444 GFP expressing-mutant strain at a final concentration of 2 × 10^6^ cell/mL in YNB media supplemented with 2% glucose and 6% ethanol. Cultures were incubated in an orbital shaker at 20°C and 150 rpm during 72 h, and 100 μL samples from each culture were extracted every 24 h. Aliquots were washed twice in PBS and stored in the same buffer. Cultures were then analyzed in a BD FACScanto II Cytometer (Biosciences, USA). Finally, the proportion of non-fluorescent/GFP-fluorescent cells was estimated. Experiments were performed in triplicate.

### Sequencing of the evolved lines and identification of mutations

DNA extraction was performed as previously described [27, 29]. Sequencing of three parallel populations at final and intermediate stages of the evolution process was performed using the Illumina HiSeq X ten platform (BGI sequencing, China). Overall, approximately 45 million reads (paired-end) were obtained for each evolved line. The raw reads were processed to remove adaptor sequences using the Fastp tool and filtered considering a 20 phred score cut-off [35]. Reads were aligned against the *S. eubayanus* CBS12357^T^ reference genome [36] using the Burrows-Wheeler Aligner [37]. Overall, 99% of the reads were aligned, obtaining a mean coverage of 980X. Genome sequences of 27 parental strains were previously sequenced [27], from which a list of SNPs that were unique for each of those sequenced strains was obtained, using a custom R script. To estimate the proportion of the parental genetic backgrounds in every evolved line, the alternative genotype coverage at each unique SNP coordinate was obtained using bcftools mpileup [38] [39]. De novo SNP calling in the evolved lines was performed using freebayes v 1.3.0 (https://github.com/ekg/freebayes). The total number of SNPs was calculated using Freebayes [40] and the effect of each SNP was predicted with SnpEff [41] and the *S. eubayanus* CBS12357^T^ reference genome [36]. Reads are available in the Biosample Database Project PRJNA666059.

### Genome reconstruction of the CLEt5.1 mutant

The genome of the CLEt5.1 mutant was reconstructed using Nanopore sequencing coupled with Illumina sequencing. Nanopore sequencing was performed using a minION system (Oxford Nanopore, Oxford, UK). For this, DNA extraction and sequencing proceeded as previously described [29]. Overall, 26.1 million reads for Illumina and 96,000 reads for Nanopore were obtained (**Table S2**). The raw fast5 files were transformed to fastq files and debarcoded using Guppy 2.3.5 [42]. Barcode and adapter sequences were trimmed using Porechop (https://github.com/rrwick/Porechop) and filtered with Filtlong (https://github.com/rrwick/Filtlong) using a Phred score of 30. Genome assembly was performed with Canu (https://github.com/marbl/canu) using default settings. Additionally, two rounds of nanopolish (https://github.com/jts/nanopolish) and pilon (https://github.com/broadinstitute/pilon) were carried out. Moreover, the raw assembly was polished using the Illumina reads filtered with a Phred score of 20 (Burrows-Wheeler Aligner). The genome assembly was annotated with the pipeline LRSDAY [43] using the *S. eubayanus* CBS12357^T^ reference genome as model for training AUGUSTUS [44], supported by the transcriptome assembly produced by TRINITY [45]. The completeness of the genome assembly was evaluated using BUSCO [46]. The assembly was compared with CBS12357^T^ using nucmer (Marçais et al, 2018) to evaluate the synteny, whilst specific structural variants (SVs) were identified using MUM&Co [47]. All the parameters of the pipeline were set up as default. The enrichment analysis of Gene Ontology (GO) terms and KEGG pathways was performed using METASCAPE [48]. The identification of transcription factor binding sites in the regulatory region 500 bp upstream of the upregulated genes of the evolved strain was performed using CiiDER [49]. Reads are available in the Biosample Database Project PRJNA666059.

### RNA-sequencing and differential expression analysis

RNA was extracted using the E.Z.N.A.^®^ Total RNA Kit I (Omega Bio-tek, USA). RNA was DNase I treated (ThermoFisher, USA) and purified using the RNeasy MinElute Cleanup Kit (Qiagen, Germany). The Illumina libraries and sequencing were performed as previously described [29] in the BGI facilities (Hong Kong, China). Briefly, RNA integrity was confirmed using a Fragment Analyzer (Agilent, USA). The RNA-seq libraries were constructed using the TruSeq RNA Sample Prep Kit v2 (Illumina, USA). The sequencing was conducted using paired-end 100-bp reads on an Illumina HiSeq X Ten in a single lane for the six samples. Reads are available in the Biosample Database Project PRJNA666059. Reads were mapped to the *S. eubayanus*CBS12357^T^ reference genome using RNAstar ver. 2.7.3 [50] and analyzed using featurecounts in R [51]. Differential expression was analyzed statistically using DESeq2 package in R [52]. Genes showing an adjusted P-value of 0.05 or less were considered as differentially expressed genes (DEGs). Analysis of GO term enrichment was performed with the R package enrichGO (https://www.rdocumentation.org/packages/clusterProfiler/versions/3.0.4/topics/enrichGO). Cytoscape was used to visualize transcription factor regulatory networks [53].

## RESULTS

### *S. eubayanus* fitness sensitivity under high ethanol conditions

We performed a parallel population assay to obtain high ethanol-tolerant *S. eubayanus* individuals (**Figure 1A**). For this, thirty *S. eubayanus* strains belonging to the PB-2 and PB-3 lineages, previously isolated in southern Chile (Villarrica, Coyhaique and Puyehue; [27]), were selected and characterized for microbial growth under different ethanol conditions. Initially, we used micro-cultures to evaluate biomass generation in 8%, 9% and 10% ethanol. Growth under these conditions showed long lag phases and low growth rates for all strains in concentrations above 9% ethanol (**Figure 1B, Table S3**). This growth was significantly lower compared to that of the L299 wine *S. cerevisiae* control strain (p-value < 0.05, ANOVA), demonstrating a greater susceptibility of *S. eubayanus* to high ethanol concentrations (**Table S3**). Furthermore, for all tested parameters, micro-culture assays demonstrated significant phenotypic differences between strains (**Figure 1B**), representing a genetically and phenotypically heterogeneous group of strains, ideal for the parallel population assay. Based on the above, we chose 9% ethanol as our selective environment for the experimental evolution procedure (from now on referred to as EtOH).

Our population assay began by mixing the thirty strains in equal proportions and subdividing them into ten mock replicates (YNB-glucose media, from now on referred to as GLU) and ten EtOH lines (**Figure 1A**). The ethanol fitness of each evolved line was evaluated at different time points during the progression of the assay (**Figure 1C**). After 260 generations (approximately six months), all GLU lines showed a significant decrease in ethanol fitness compared to the ancestral culture (p-value < 0.05, ANOVA, **Figure 1C**). In contrast, the EtOH-evolved lines showed higher maximum growth rates (μmax) in ethanol compared to the original mixed-culture, attaining a 60% greater μmax (p-value < 0.05, ANOVA). These differences were not observed in glucose micro-cultures (**Figure 1D**). Thus, demonstrating that the evolved lines performed better in their selective environment compared to the control condition. Interestingly, we did not detect major adverse phenotypic effects in beer wort, suggesting a low accumulation of detrimental mutations (**Table S3b**).

### Genome sequencing reveals consistent strain selection in parallel populations

Three GLU (GLU-1, 7 and 10) and three EtOH lines (EtOH-2, 5 and 6) were sequenced at the end of the experiment to identify the genomic changes and the pervasiveness of the different genetic backgrounds across the assay. Interestingly, all the EtOH sequenced lines showed a sustained prevalence of strain CL248.1 (belonging to PB-2 and isolated in northern Patagonia), reaching over 95% of the population’s allele frequency by the end of the experimental evolution assay (**Figure 2A)**. That being said, CL248.1 did not show the highest growth rare (μmax) under ethanol 9% of the *S. eubayanus* strains considered in this study, suggesting that selection did not occur solely due to ethanol tolerance (**Table S3**). In contrast, we did not observe a consistent selection in the GLU lines, where different genetic backgrounds were found depending on the evolved line (**Figure 2A**). These results likely suggest a milder and different selection pressure in yeast when glucose is used as a selection regime, and a particular competitive fitness advantage of CL248.1 solely under EtOH selection, demonstrating a convergent phenomenon when ethanol and biotic stress are applied together.

**Figure 2.**
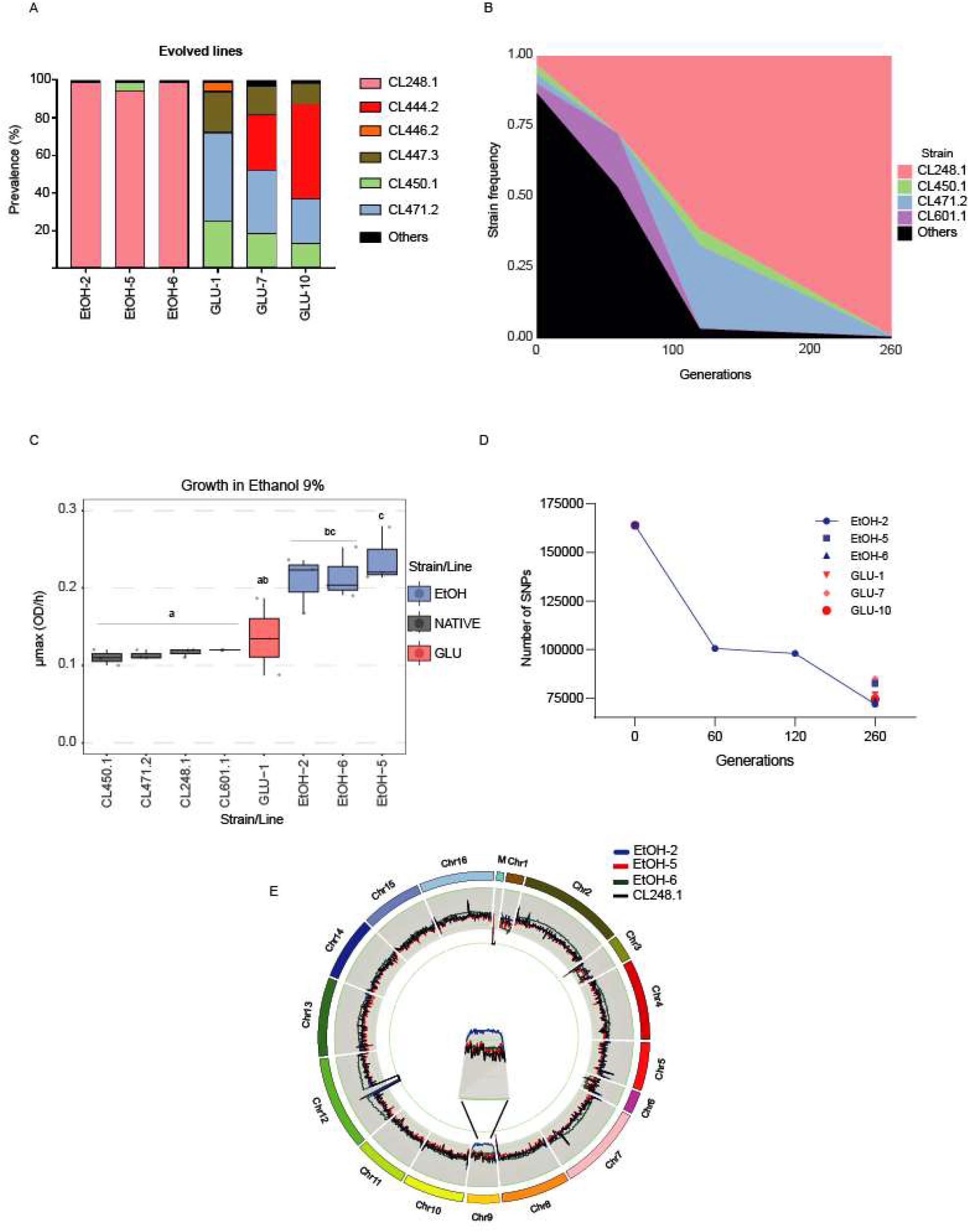
Genomic and phenotypic changes in the evolved lines. (A) The presence of the prevalent genetic backgrounds in three glucose (GLU) and ethanol (EtOH) evolved lines. (B) Prevalence (frequency) of the most prominent genetic backgrounds during the evolution of line EtOH-2. (C) Ethanol 9% growth rates for the most representative parental strains and evolved lines. (D) Total number of SNPs relative to the CBS12357 reference genome at the beginning and end of the evolution assay for EtOH-5, 6 and GLU-1, 7 and 10 lines. In addition, the number of SNPs during the evolution assay is shown for EtOH-2. (E) Chromosome number estimation across EtOH lines. Only the EtOH-2 line showed an aneuploidy by the end of the evolution assay (chromosome 9).

Line EtOH-2 was sequenced at different time points (0, 60, 120 and 260 generations) to identify the genotypic course of the assay and additional genotypes under selection (**Figure 2B**). We observed a predominance of CL248.1 and CL601.1 genotypes after 60 generations, demonstrating a competitive displacement of CL248.1 in the culture, together with higher fitness over the other genetic backgrounds (**Figure 2B**). Interestingly, after 120 generations, four genotypes monopolized the culture, representing 96.6% of the EtOH-2 line. Nevertheless, none of these parental genotypes showed high ethanol growth rates compared to the evolved lines (**Figure 2C**). A second genotype, CL471.1 reached significant frequencies (maxima 29.3%) during intermediate periods of the evolution assay. However, it was almost absent by the end of the experiment, being detected at a frequency of just 0.15% in the final population. Moreover, over time, we calculated the total number of SNPs in evolved lines against the reference strain CBS12357^T^. We found a decrease in the number of SNPs over time across all lines relative to the ancestral culture, particularly in EtOH-2, which exhibited the greatest decay compared to other lines (**Figure 2D**).

To identify de novo genetic variants with a potential effect on ethanol tolerance, we used the EtOH-2 line and compared polymorphisms (SNPs and short INDELs identified using freebayes) before and after selection. We chose this line because it showed the highest homology to a single genetic background (CL248.1), allowing the identification of novel genetic variants over the raw population’s genetic variation. In this way, we arbitrarily selected for polymorphisms with a putative moderate/high impact on the gene function and found 34 impacted genes under these criteria (**Table S4**). Among others, we found mutations in genes such as *YPS6* and *IMA1,* encoding for a putative GPI-anchored aspartic protease [54] and a isomaltase [55], respectively. We also found a single aneuploidy in the EtOH-2 line in chromosome IX, where an extra copy was found (**Figure 2E).**Altogether, our results demonstrate how ethanol promotes a significant decrease in genetic variability due to genotype selection coupled with the emergence of new adaptive mutations vital for ethanol survival in biological processes such as stress damage and sugar metabolism.

### Ethanol-evolved individuals have greater fermentation capacity and maltotriose consumption

We determined the fitness cost of ethanol adaptation in 24 different environmental conditions for those EtOH adapted individuals isolated after 260 generations of selection. For this, we randomly isolated two clones from each EtOH line and estimated growth rates in micro-cultures considering diverse phenotypic growth conditions, including high temperature, different carbon sources, and oxidative and osmotic stress (**Table S5**). To control for adaptive mutations in YNB laboratory media, we also isolated two colonies from three GLU lines. In general, individuals from EtOH-evolved lines showed higher μmax in ethanol (**Figure 3A**), and also for a greater number of conditions, compared to GLU-evolved individuals and the ancestral culture (p-value < 0.05, ANOVA, **Table S5**). These conditions included greater growth rates in sources such as glucose, maltose and fructose, together with resistance to oxidative (H2O2) and osmotic stresses (sorbitol 20%) (**Figure 3B**), suggesting that selection improved general stress tolerance in these evolved strains. Interestingly, we found that one EtOH evolved individual (CLEt5.1, isolate n°1 from the EtOH-5 line) exhibited greater ethanol tolerance, but a lower growth rate under high temperature (34°C) and an ionic detergent (SDS 0.001%, **Figure 3B**), indicating the existence of a trade-off.

**Figure 3.**
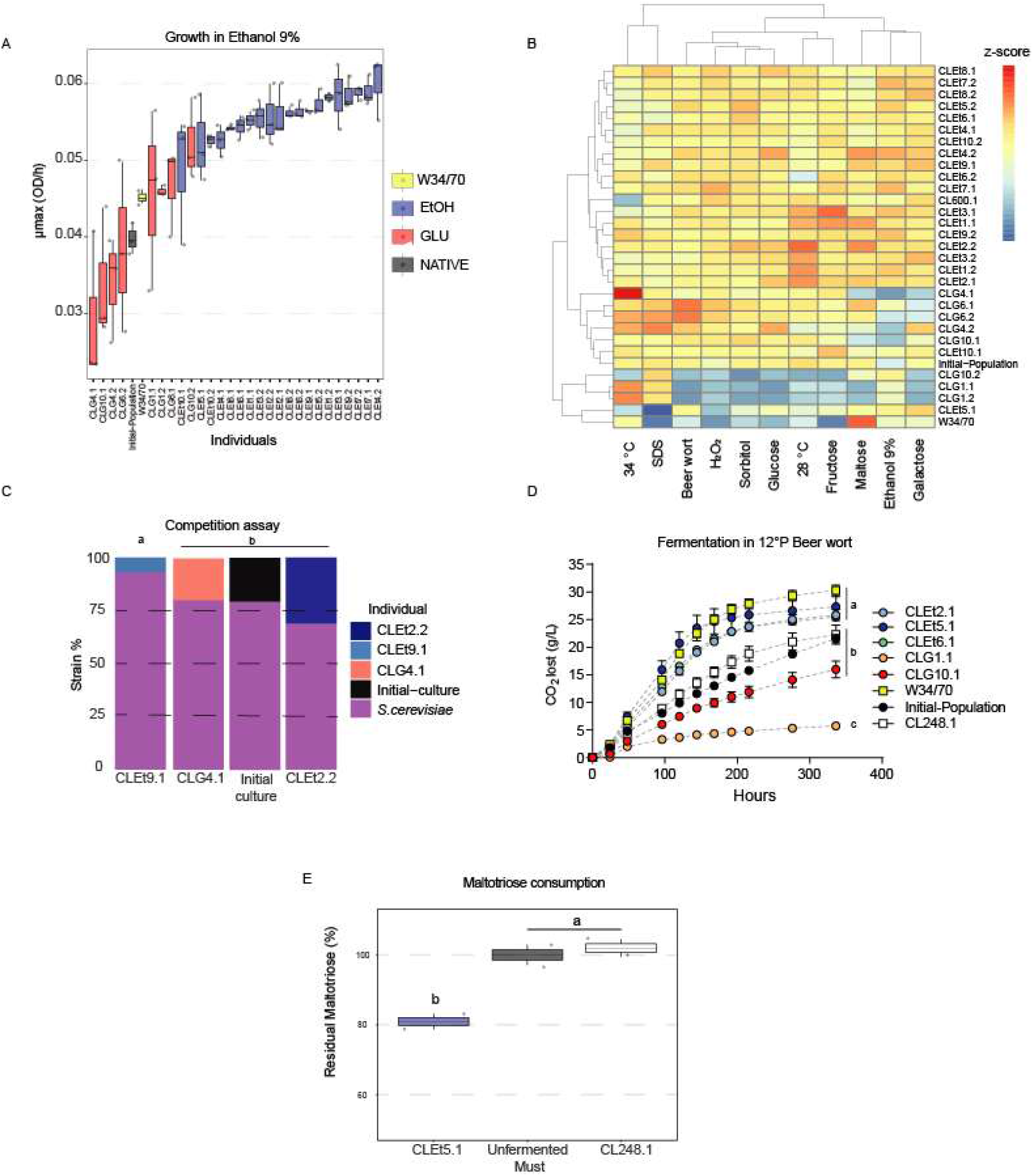
Phenotypic profiling of evolved individuals. (A) Growth rates under YNB-glucose-ethanol 9% of different evolved individuals. (B) Phenotypic heatmap based on micro-culture growth rates of EtOH and GLU-evolved individuals, evaluated in 11 different conditions. (C) The evolved strains were challenged using a GFP-mutant *S. cerevisiae* in YNB media supplemented with ethanol at 9%. The strain frequency of the evolved individuals was evaluated using flow cytometry. (D) Fermentation in 12 °P beer wort of different evolved individuals. The fermentative capacity was estimated from the CO_2_ lost at different time-points. The statistical differences were calculated after 216 h of fermentation using ANOVA. (E) Maltotriose consumption in YNB maltotriose 2% micro-cultures.

To determine the relative fitness of EtOH and GLU evolved individuals, we carried out a competition assay in YNB-glucose supplemented with 6% ethanol, against a recombinant *S. cerevisiae* that constitutively expresses GFP (**Figure S1**). We observed that all tested strains were unable to outcompete *S. cerevisiae*; however, significant differences were found in the final proportion of the tested strains at the end of the experiment (*p*-value < 0.05, ANOVA). For example, strain CLEt9.1 was almost absent at the end of the competition assay (relative frequency < 0.1), while CLEt2.2 was found to represent 31% of the cells quantified in the final culture (**Figure 3C**). These results demonstrate fitness differences between EtOH and GLU isolated individuals.

Additionally, we evaluated the fermentative capacity in small-scale lager wort fermentations at low temperature (12 °C) of three EtOH and two GLU evolved individuals. The selected strains were monitored for 15 days, and their fermentative capacity was estimated by measuring CO_2_ loss and sugar consumption throughout the fermentative process (**Figure 3D**). Surprisingly, all the EtOH-evolved individuals showed a similar fermentative profile compared to the commercial strain, where no significant differences were found in terms of total CO_2_ loss (p-value < 0.05, ANOVA). Furthermore, the best-evolved isolate (CLEt5.1) showed a 22.6% increase in loss of CO_2_ compared with the ancestral culture after 14 days of fermentation (**Figure 3D** and **Figure S2A**, p-value > 0.05, ANOVA), and also exceeded the fermentative performance, in terms of fermentation rate, of its parental genetic background CL248.1 (**Figure 3D, S2B, and S2C**). Moreover, sugar consumption differed between the W34/70 commercial strain and the evolved individuals. Although the isolates were able to consume all the glucose, maltose, and fructose found in the wort (**Figure S2D**), no maltotriose consumption was observed (p-value < 0.05, ANOVA, **Table S6**) in the evolved strains. We only detected maltotriose consumption under fermentation conditions in the lager commercial strain, in agreement with the inability of *S. eubayanus* to use this carbon source (**Figure S2D**, [23]). To further analyze maltotriose consumption, we quantified the remaining maltotriose concentration after a 5-day incubation period of the evolved individuals in YNB synthetic media supplemented with 2% maltotriose as the sole carbon source (**Figure 3E**). Interestingly, we detected 19.1% maltotriose consumption in the evolved strain CLEt5.1, while no consumption was found in CL248.1 (**Figure 3E**). These results suggest genomic and molecular changes leading to maltotriose metabolization in this genetic background that only arise when maltotriose is used as the sole carbon source.

### Identification of *de novo* genetic variants in the EtOH evolved strain CLEt5.1

The genome of the EtOH-evolved individual CLEt5.1 was sequenced by coupling Nanopore and Illumina technologies to elucidate the genetic origin of the phenotypic changes acquired through the evolution process. We obtained a high-quality assembly and identified 5,946 genes in the final genome annotation, organized in 37 scaffolds (**Figure 4, Table S2 and Table S7a)**. The completeness analysis using BUSCO showed that the *de novo* assembly contained almost all the expected set of genes for a member of the *Saccharomyces* genus (97.5%). By comparing the scaffolds of the assembly against the CBS12357^T^ reference genome, high synteny between genomes was observed, except for an evident translocation between chromosomes IV-R and XVI-L (**Figure 4**). Therefore, we proceeded to identify structural variants between CLEt5.1 and its parental background (CL248.1) using MUM&Co [47]. In this way, we identified 100 structural variants (Deletions: 47, Insertions: 41, Duplications: 10, Inversions: 0 and Translocations: 2, **Table S7b**), primordially INDELs and confirming the translocation between chromosomes IV-R and XVI-L of 980 kb. Additionally, we found a 47 kb deletion in chromosome XII, and two 24 kb and 39 kb duplications in chromosomes VII and IV, respectively. Among the genes present in the chromosome VII duplication, we found *VID30,* which is involved in the regulation of carbohydrate metabolism and the balance of nitrogen metabolism towards glutamate production, and *HAP2,* a transcription factor which is predicted to regulate many of the proteins induced during the diauxic shift [56](**Table S7c**). SNP calling using freebayes detected 1,006 high quality SNPs.

**Figure 4.**
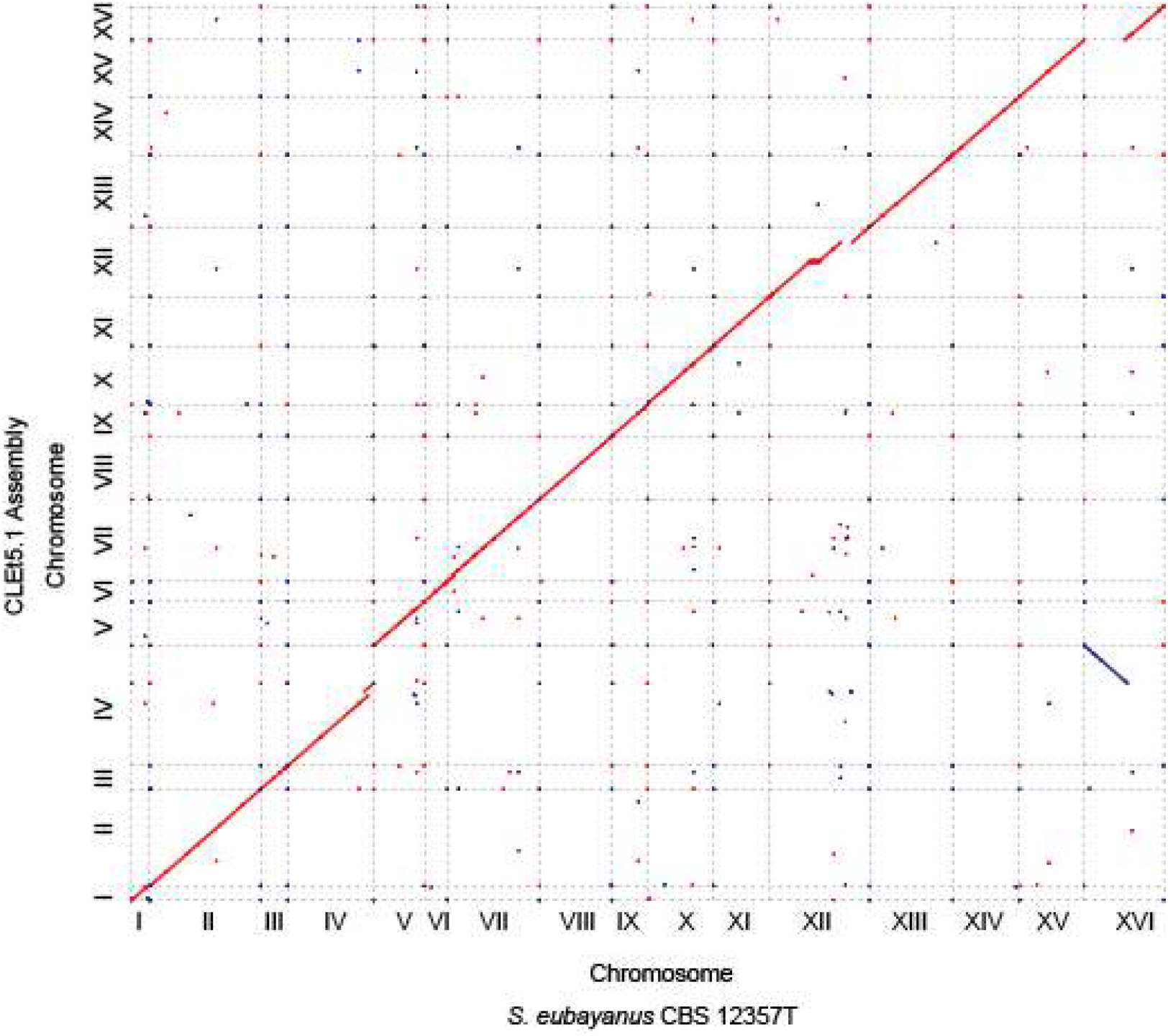
Genome synteny analysis of the EtOH evolved CLEt5.1 strain. Dot plot representation of DNA sequence identity between the *S. eubayanus* CBS12357^T^ strain and the EtOH evolved CLEt5.1 strain. A single translocation was found between chromosome IV and XVI.

To better understand the molecular basis of ethanol adaptation, we searched for polymorphisms across the CLEt5.1 genome that could generate moderate or high impact mutations on the gene function (based on snpeff predictions). We found 11 genes with significant polymorphisms between CLEt5.1 and the native CL248.1 strain (**Table S7d**). For example, we found a missense variant in *PUT4,* which encodes for a proline permease essential in proline assimilation during fermentation [57]. Similarly, we found a frameshift in *IRA2,* which encodes for a GTPase-activating protein, and previously related to high-temperature fermentation [58] and low glucose-growth defect rescue [59]. These results demonstrate that this relatively short period of ethanol adaptation promoted punctual, small and large rearrangements, which, taken together may be responsible for the phenotypic differences between the CLEt5.1 and CL248.1 strains.

### Transcriptome and organoleptic analysis of the CLEt5.1 evolved strain under beer fermentation

To determine the impact of genetic changes in metabolic processes during wort fermentation in EtOH adapted individuals, we used a transcriptome approach. This allowed us to identify differentially expressed genes (DEGs) between the CLEt5.1 and the CL248.1 parental strain after 24 h of fermentation in a 1.5 L fermenter. Overall, we observed 92 DEGs (Fold change > 0.7 and FDR < 0.05, **Figure 5A** and **Table S8**), of which 59 and 33 were up- and down-regulated in the CLEt5.1 strain, respectively. Enrichment analysis of GO terms in up-regulated genes revealed that diverse biological and molecular pathways, including sulfur compounds, methionine metabolism, and several cellular amino acid metabolic processes were enriched in the evolved strain (**Table S8**). In contrast, down-regulated genes were significantly enriched in alpha-amino acid metabolism and pheromone response metabolism, together with cofactor and vitamin binding molecular functions (**Table S8**). Similarly, KEGG enrichment analysis highlighted that genes within several pathways were differentially expressed between genotypes. For example, assimilatory sulfate reduction, cysteine and methionine metabolism, seleno-compound metabolism and biosynthesis of antibiotics pathways were enriched in the up-regulated genes set (**Table S8**). In contrast, we found a significant enrichment of the amino acid biosynthesis pathway among down-regulated genes (p-value < 0.01, hypergeometric test). Interestingly, these two analyses highlight that several DEGs were related to nitrogen and amino acid uptake, stress tolerance, and faster diauxic shift, suggesting that nitrogen uptake and a rapid stress response play essential roles during fermentation in this evolved strain.

**Figure 5.**
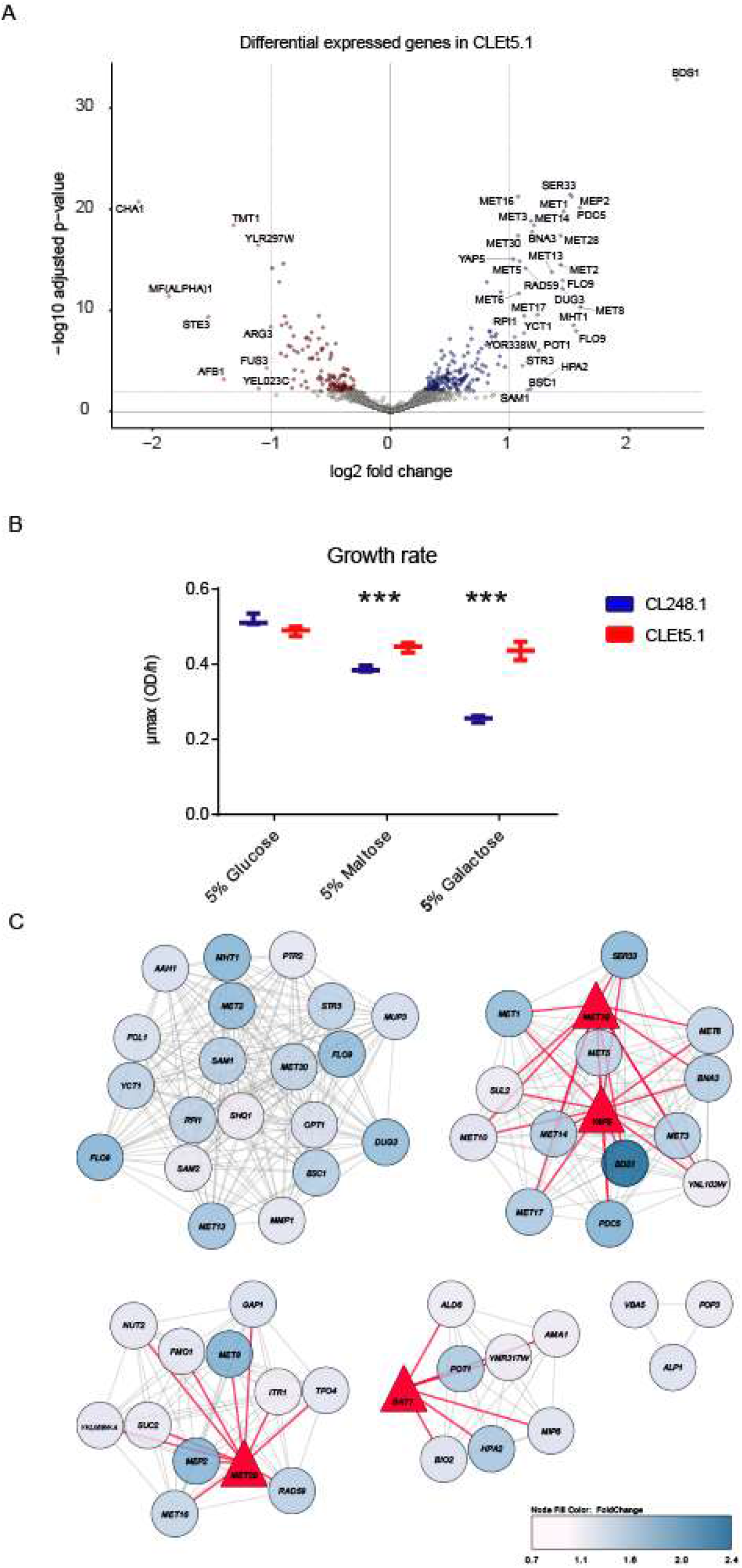
Differential gene expression analysis between the EtOH evolved CLEt5.1 strain and its native parental strain under beer wort fermentation conditions. The transcriptome of the CLEt5.1 EtOH evolved strain was evaluated and compared against the CL248.1 native strain under beer wort fermentation conditions. (A) The volcano plot depicts differentially expressed genes between CLEt5.1 and CL248.1 (B) Relative growth rates of CLEt5.1 and CL248.1 strains shifted from two 24 hours 5% glucose pre-cultures to 5% maltose and 5% galactose media. (C) Network analysis in upregulated genes in CLEt5.1 depicting the most relevant hubs differently regulating genes between CLEt5.1 and CL248.1. Transcription factors are shown in red triangles, while TF-gene connections are shown in red lines.

To evaluate the fast diauxic shift and the capacity of these two strains to switch from glucose to other disaccharides, we estimated their growth capacity under maltose and galactose after two 24 h pre-cultures in 5% glucose. In agreement with our transcriptome results, the evolved strain showed a significantly greater growth rate compared to CL248.1 under 6% maltose and 6% galactose concentrations after long glucose incubation periods (**Figure 5B**).

Additionally, to identify possible common regulatory elements of the up-regulated genes, we analyzed their promoter sequences (500 bp upstream of the transcription start site), and found a significant enrichment of transcription factor binding sites (p-value < 0.05, Fisher’s exact test) for transcription co-activators of the Cbf1-Met4-Met28p complex (methionine metabolism), Dal80p and Uga3p (activators of nitrogen metabolism), Tye7p (glycolytic genes activator) and Sfl1p (repression of flocculation-related genes, and activation of stress responsive genes, **Table S9**). Additionally, we used Cytoscape to visualize the resulting network predicting regulatory interactions from the set of upregulating genes (**Figure 5C**). According to our network model, we found four transcription factors: Met28p, Met32p, Gatp and Yap5p modulating the expression of these up-regulated genes in CLEt5.1. Interestingly, Yap5p is known to be involved in the diauxic shift [60]. These results highlight a transcriptional rewiring in CLEt5.1 for genes related with nutrient acquisition, stress tolerance and methionine metabolism during the evolution of tolerance to fermentation stress.

During the fermentation process, we subjectively perceived that the organoleptic properties of the beers produced by the evolved strain differed from those of the parental native strain. Therefore, to determine how the transcriptional rewiring and genomic changes impacted the production of volatile compounds and the beer profile in the CLEt5.1 evolved strain, we quantified volatile compound production using HS-SPME-GC/MS at the end of fermentation (day 15). As expected, we found significant differences in the composition of volatile compounds produced in beer between the evolved and parental strains (p-value < 0.05, paired t-test, **Figure 6A, Table S10**). In general, the evolved clone showed lower levels of ester compounds, such as isoamyl acetate and ethyl octanoate (p-value < 0.05, ANOVA). Additionally, we detected high levels of benzaldehyde 4-methyl (aromatic aldehyde) and ethyl hexadecanoate in the evolved strain compared to the native genetic background, which could confer a fruity aroma to the beer similar to those found in lager beers. The most interesting differences were found in terms of off-flavors. We detected a significantly lower production of 2-Methoxy-4-vinylphenol (4-vinyl guaiacol) in the evolved strain, likely reducing its clove-like flavor, which is typically found in fermented beverages by wild strains (p-value < 0.05, ANOVA**, Figure 6A**). Interestingly, we did not find mutations in the *FDC1* and *PAD1* coding regions, or a significant difference in gene expression for *FDC1* (log2FC = −0.038, p-value adjusted = 0.838) and *PAD1* (log2FC = −0.0095, p-value adjusted = 0.965) between both strains. However, a series of mutations in the regulatory regions of both genes were found in CLEt5.1, which could alter expression levels in later fermentation stages. These results suggest that the evolution process significantly impacted the volatile compound profile of beers produced by CLEt5.1, emulating the domestication process that modified several commercial yeasts.

**Figure 6.**
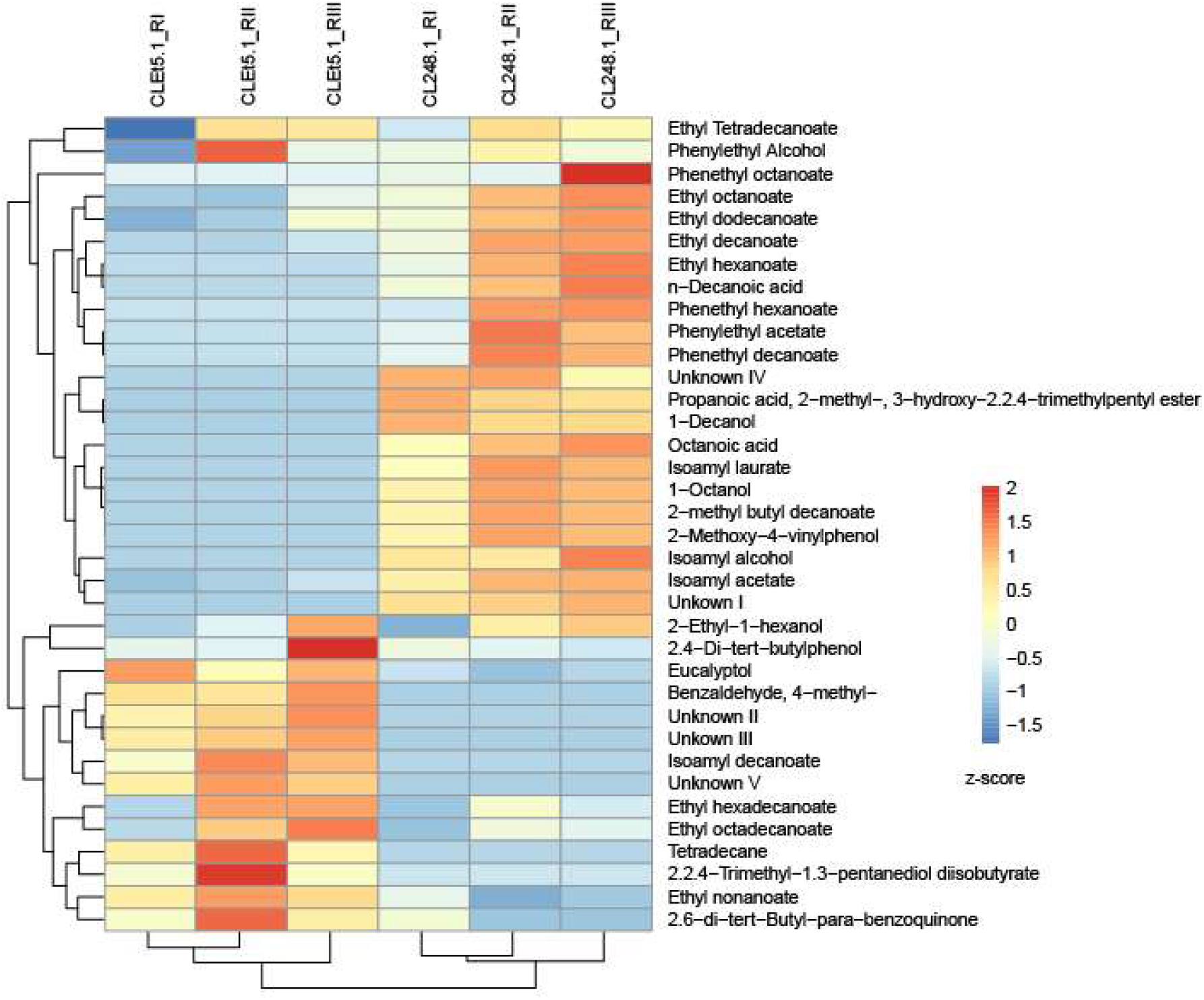
Volatile compound production on beer wort. . The final beer from EtOH evolved CLEt5.1 and its parental strain CL248.1 was analyzed using HS-SPME-GC/MS. The relative abundance of each compound detected was evaluated and a heatmap was constructed. The compounds were grouped in accordance to their relative abundance.

## DISCUSSION

Human-driven selection associated with yeast domestication in fermentative environments has been extensively reported in *S. cerevisiae* and related hybrids [14, 15]. However, the genetic basis and molecular changes in other *Saccharomyces* genomes associated with alcoholic beverages is still unclear. In our study, we have reconstructed the putative domestication history of the yeast *S. eubayanus* under biotic and abiotic stresses, using a panmictic founding population that simulated the natural process of adaptive evolution, and using an ethanol environment as the selective agent. We used dozens of wild genotypes in a single culture, in order to replicate the natural genetic variability of these organisms. We observed that a single genetic background, CL248.1, systematically outcompetes the others, acquiring *de novo* mutations and improving basal ethanol tolerance. Interestingly, the time-course of this competitive displacement was complex, involving genotype selection and innovations throughout the assay (key adaptive mutations) that were constantly replaced by others during the “fast-motion” evolution time-course. Thus, the evolved lineages derived from our founding genetic background exhibited higher ethanol growth rates compared to their ancestors, demonstrating a rapid response to selection, and so adapted successfully to their new environment. However, CL248.1 was not the best ethanol-tolerant strain, suggesting that pre-existing variants, together with *de novo* mutations, combined to positively-affect fitness in this strain. In this sense, it has been demonstrated that pre-existing and *de novo* genetic variants can both drive long term adaptation to environmental changes in yeast [61]. This indicates that not only a fitness advantage related to a given environmental selection pressure is essential for directional selection to occur in populations [62], but also that a combination of standing genetic variation with some genomic plasticity for beneficial mutations are essential [63]. In this way, the success of an individual is established in such a competitive environment [64]. Our results show that both pre-existing genetic variation and *de novo* mutations of a range of effects were important in explaining rapid evolution in this ecological context [65, 66]. Importantly, the *Saccharomyces* “make-accumulate-consume (ethanol)” life strategy is fundamental for withstanding the antimicrobial effects of ethanol in a complex population [67, 68]. Thanks to this, multiple *Saccharomyces* genotypes were selected, domesticated, and used over centuries in the beer industries, including the *S. pastorianus* hybrid [14, 15].

Domestication signatures in yeast, as a result of the human-domestication syndrome, included genomic changes in the *S. cerevisiae* and *S. eubayanus* genomic portions leading to faster fermentation rates under low temperatures, a more moderate organoleptic complexity, and the absence of off-flavors in beers [14, 20]. Under the premise that evolutionary experiments can lead to unexpected and somewhat counterintuitive results [69], we evaluated the beer fermentation performance of *S. eubayanus* evolved individuals. Interestingly, evolved individuals exhibited a similar fermentation performance compared to lager yeast, suggesting in turn that ethanol, together with competitive displacement, could be the leading drivers of yeast domestication in brewing environments. This persistent directional selection involved correlated selection of other traits, such as osmotic stress tolerance and efficient nitrogen uptake [70]. In general, domesticated fungi used in fermented foods exhibit genomic rearrangements, fewer spores and produce desirable volatile compounds [9]. These domestication signatures have been reported in other systems, such as *Aspergillus* and *Penicillium,* where a transition to environments rich in carbon and nitrogen sources led to extensive metabolism remodeling when used to produce cheese [8, 9].

Ethanol-evolved individuals presented a series of genomic changes related to yeast domestication, such as aneuploidy and chromosomal rearrangements [16]. Furthermore, signatures of trait domestication are evident in evolved individuals showing improved stress resistance, fast fermentation rates, lower organoleptic complexity and a lower production of phenolic off-flavors [14]. *S. cerevisiae* beer strains are characterized by strong domestication signatures in their genomes, including polyploidies, the decay of sexual reproduction, and maltotriose consumption [16]. Interestingly, one of our strains was able to consume maltotriose, which is another key domestication hallmark. In terms of the molecular mechanisms that explain their increased fermentative capacity, we observed that some stress response genes were either mutated or up-regulated in the ethanol-evolved line compared to its parental genetic background. In this way, the mutations and genomic rearrangements found in the CLEt5.1 evolved individual could explain the transcriptional rewiring and improved fermentative profile. Indeed, ethanol exposure leads to the recruitment of error-prone DNA polymerases, causing DNA replication stress and increased mutation rates [71]. Accordingly, we found that *RAD59* (involved in DNA double-strand break repair) was overexpressed in the evolved strain CLEt5.1, likely indicative of a mechanism that counteracts the mutagenic effect of ethanol [72]. Other overexpressed genes could also be directly related to an increased fermentative capacity, such as *SUC2, YAP5* and *MET*, which could promote glucose uptake, a dynamic diauxic shift, and the accumulation of S-Adenosylmethionine, respectively [73, 74 2013]. In this context, genomic rearrangements, such as the duplication found in chromosome VII containing *HAP2,* which is involved in promoting the diauxic shift, are in agreement with these findings. Furthermore, previous reports in lager yeast demonstrated that the accumulation and exogenous supplementation of S-Adenosylmethionine promotes an increase in the fermentative capacity of yeast under high-gravity wort [75].

### Concluding remarks

In summary, the results found in our study could be applied to determine the domestication dynamics of the *S. eubayanus* genomic portion in the lager strain, given the occurrence of similar desirable traits for beer. Based on multiple analyses, we provide evidence of the intermediate evolutionary changes in *S. eubayanus,* which have direct implications in the generation of novel yeasts for the industry. In this way, genomic changes promote a transcriptional rewiring that induces a favorable response in a fermentative environment. For the first time, these findings provide novel insights into the genomic and phenomic changes in wild *S. eubayanus* leading to faster wort fermentation rates and desirable organoleptic complexity, demonstrating its broad feasible use in the beer industry.

## ACKNOWLEDGEMENTS

This research is supported to FC by Comisión Nacional de Investigación Científica y Tecnológica CONICYT FONDECYT [1180161] and Millennium Institute for Integrative Biology (iBio). WM is supported by CONICYT FONDECYT [grant 3190532]. CV is supported by CONICYT FONDECYT [grant 3170404]. JM is supported by ANID FONDECYT POSTDOCTORADO [grant 3200545]. RN is supported by FIC ‘Transferencia Levaduras Nativas para Cerveza Artesanal’ and Fondecyt grant [1180917]. We thank Michael Handford (Universidad de Chile) for language support.

## SUPPLEMENTARY MATERIAL

### SUPPLEMENTARY FIGURES

Figure S1. **Competition assay of evolved individual in ethanol 9%.**

Figure S2. **Fermentative capacity of the evolved individuals**. (A) The fermentative capacity is indicated as a percentage of the capacity of the *S. pastorianus* control strain (W34/70) at 7 days. The fermentative capacity was estimated from the loss of CO_2_ over time. All assays were performed in triplicate. (B) The fermentative capacity was also determined at 14 days. (C) The velocity of the fermentation was estimated and (D) the residual sugars and metabolites in the wort were evaluated using HPLC.

### SUPPLEMENTARY TABLE LEGENDS

**Table S1. Native *S. eubayanus* strains used in the experimental evolution assay**. The strain ID and the location of isolation site are indicated.

**Table S2. Bioinformatics Summary statistics**

**Table S3. Growth kinetic parameters in glucose and ethanol of the native parental strains used for the ancestral culture**. Growth parameters μmax (OD/hr), OD max (OD) and lag phase (1/hr).

**Table S4. SNPeffect analysis of the novel polymorphisms in EtOH-2.** Snpeffect analysis of the novel/fixed polymorphisms in EtOH-2 after 260 generations

**Table S5. Phenotype data of evolved individuals.** The data shows the average μmax across three replicates and the standard deviation (SD) for diverse growth conditions, including high temperature (28°C and 34°C), different carbon sources (glucose, fructose, maltose, galactose, xylose), and oxidative (ethanol 9%, 3 mM H2O2) and osmotic stress (beer wort, SDS 0.001%, Sorbitol 20%).

**Table S6. Sugar consumption and metabolite production of the evolved individuals from fermentations in beer wort**. Sugar consumption (g/L) and metabolite production (g/L) are informed.

**Table S7. Structural variants identified in CLEt5.1 using MUM&Co.** A. CLEt5.1 genome assembly and annotation statistics. The genome assembly of CLEt5.1 using Nanopore and Illumina sequencing technology was used to calculate several assembly statistics. B. All structural variants. C. Duplicated genes present in the chromosome IV - chromosome XVI duplication in CLEt5.1. D. High/moderate SNPeff prediction of SNPs and short INDELs in CLEt5.1

**Table S8. Differential gene expression between CL248.1 and CLEt5.1 under beer wort.** A. Gene expression results. B. Upregulated and C. Downregulated genes in CLEt5.1. R1, R2 and R3 represent the three biological replicates for each genotype.

**Table S9. Enrichment analysis of Transcription Factor binding sites in regulatory regions of upregulated genes using CiiDER.**

**Table S10. Volatile compound production in CL248.1 and CLEt5.1 in beer wort.**

